# Characterization of the BRAF interactome identifies BRAF^V600E^<=>TP53 interaction in melanoma

**DOI:** 10.1101/2025.06.20.660711

**Authors:** Kayla T. O’Toole, Adamaris Martinez, Brandon Murphy, Anastasia Proveyeka, Gabriela Fort, Fatima Al-Sudani, Sanjana Boggaram, Elliott L. Paine, Deevya Baral, Joshua L. Andersen, Gennie Parkman, Eric L. Snyder, Robert Judson-Torres, Martin McMahon

## Abstract

Melanoma is a highly aggressive and frequently metastatic cancer with its incidence reported to be on the rise. Although most oncogenic drivers in melanoma converge on activation of the RAS>RAF>MEK>ERK MAPK signaling pathway, not all MAPK-activating mutations are recurrently observed in this disease, suggesting a unique functional role for BRAF^V600E^, which is present in ∼50% of all melanoma cases. However, the prevalence of BRAF^V600E^ alterations over other known MAPK-promoting oncoproteins raises questions regarding whether BRAF^V600E^ possesses additional functions outside of MAPK pathway activation. Thus, we performed TurboID to differentiate the interactome between wild-type BRAF and BRAF^V600E^. We identified novel interacting partners of normal vs. BRAF^V600E^, most strikingly being the tumor suppressor TP53. While TP53 is commonly altered or lost across many malignancies, it is notable that TP53 alterations are rare in melanoma. Our studies suggest that BRAF^V600E^ can interact with and inactivate TP53, thus providing potential mechanistic explanation as to why TP53 inactivation or loss is infrequent in BRAF^V600E^-driven melanoma.

## INTRODUCTION

Mutationally-activated BRAF^V600E^ is a driver of several human cancers, including melanoma, colon, lung, and thyroid cancers. The most common mutational alteration of BRAF, *BRAF^T1799A^*, encodes the BRAF^V600E^ oncoprotein kinase which exhibits constitutive kinase activity compared to normal BRAF. Multiple ATP-competitive inhibitors have been FDA approved to target BRAF^V600E^including vemurafenib, dabrafenib, and encorafenib^1–4^. Despite the initial success of these treatment strategies, the problem of drug resistance, either primary or acquired, remains a major therapeutic obstacle to the depth and durability of patient responses^5–9^, highlighting the need to identify new therapeutic strategies.

BRAF^V600E^ is frequently referred to as a “monomer,” as it can signal independent of RAS-GTP and other RAF kinases^8,10^. Nevertheless, BRAF^V600E^ must still form complexes with other proteins such as 14-3-3, KSR, and MEK to propagate signaling^11,12^. This prompted us to investigate if BRAF^V600E^ may be able to interact with proteins unavailable to normal BRAF proteins. Additionally, protein-protein interactions control cellular functions and are drivers of cancer initiation and progression. As such, elucidating protein-protein networks provides fundamental insight into disease mechanisms and may be useful for identification of novel treatment strategies. To date, the majority of previous studies investigating the BRAF interactome have utilized conventional techniques such as IP-MS which fail to identify low-affinity interactors, including known interactors such as MEK1/2^13–18^. Here, we leverage the proximity labeling technique, TurboID^19^, to map the interactomes of both wild-type and oncogenic BRAF and uncover previously unknown BRAF protein-protein interactions. We identified over 1,300 potential BRAF^V600E^-interactors, ∼200 of which were unique compared to normal BRAF. Perhaps surprisingly, the tumor suppressor, TP53, was among enriched interactors with BRAF^V600E^.

The tumor suppressor protein TP53 plays a crucial role in maintaining genomic stability and regulating the cell cycle, apoptosis, and DNA repair. In its canonical role, TP53 acts as a “guardian of the genome”, preventing the accumulation of mutations that could lead to cancer development^20–22^. Activation of TP53 is typically triggered by signaling pathways induced by cellular stresses such as DNA damage or oncogene activation^23–25^. For this reason, TP53 is often inactivated in cancer either through point mutations or genetic deletion. However, melanoma is known to have a relatively low frequency of TP53 mutations, estimated to occur in only about 17% of cases^26^. Interestingly, studies have shown that when TP53 mutations do occur in melanoma, they often arise after the tumor’s initial development as an acquired resistance mechanism, rather than driving melanomagenesis^27–30^.

Despite the notable absence of TP53 alterations in melanoma, TP53 function can often be inhibited through alternative mechanisms, such as alterations in upstream signaling pathways, protein-protein interactions, or epigenetic modifications^31–36^. In melanoma, amplification of *MDM2* or elevated expression of *MDM4* has been shown to inactivate TP53^37–39^. However, alterations in TP53 or its subsequent regulatory proteins is notably absent in several of our BRAF^V600E^-driven cell lines, potentially suggesting alternative mechanisms of inactivation in BRAF-driven melanomas. Importantly, Mo et al., described the concept of neomorphic protein<=>protein interactions, demonstrating that mutant proteins can acquire distinct and novel binding partners that differ from their normal counterparts. In particular, their work suggested that BRAF^V600E^ can interact with a multiplicity of new binding partners compared to normal BRAF, one of which was TP53^15^. Building upon this foundation, our study not only confirms this putative interaction between BRAF^V600E^ and TP53 but also provides new insights by revealing that BRAF^V600E^ expression alone is sufficient to induce a redistribution of TP53 within the cell and attenuate TP53 transcriptional activity, even following ultraviolet (UV) irradiation or inhibition of MDM2. In addition, we demonstrate that the DNA-binding domain of TP53 is critical for this interaction with both normal and BRAF^V600E^. Our results suggest a mechanistic basis by which BRAF^V600E^ may contribute to tumor progression through modulation of TP53 function. Further, the results presented here provide a potential explanation of why TP53 inactivation or loss rarely co-occurs with the expression of the BRAF^V600E^ oncoprotein kinase.

## RESULTS

### Normal and oncogenic BRAF have unique protein-protein interactions

To understand how the V600E substitution alters BRAF function, we utilized an unbiased, BirA proximity-dependent biotin tagging identification screen (TurboID). This technique allows for the detection of weak or transient interactors that are frequently undetected using immunoprecipitation techniques and allows for a more comprehensive investigation of the BRAF interactome^19,40^. The BirA biotin ligase was fused to the C-terminus of either normal full-length human BRAF or BRAF^V600E^ ^41^ with a flexible linker to allow for biotinylation of proteins within a ∼20nm radius (Figure 1A and Supplemental Figure 1A). HEK293 cells were transduced with these doxycycline-inducible Turbo-ID constructs and immunofluorescence microscopy confirmed the TurboID construct did not impact BRAF subcellular localization, consistent with previous publications^42^, and confirmed that these constructs biotinylate proximal proteins using immunoblotting and a fluorescent neutravidin stain (Supplemental Figures 1B and 1C). Additionally, we confirmed that our TurboID constructs are induced by doxycycline treatment and immunoblotting also confirms increased intracellular biotinylation with construct expression (Supplemental Figure 1D). Lastly, we confirmed the TurboID constructs were resistant to siRNAs targeting endogenous BRAF, which were utilized in order to maximize the BRAF-TurboID signal. Immunoblotting demonstrated elevated phosphorylated ERK (pT202/Y204) with the expression of the BRAF-TurboID constructs, characteristic of activated RAF signaling (Supplemental Figures 1E and 1F).

**Figure 1:**
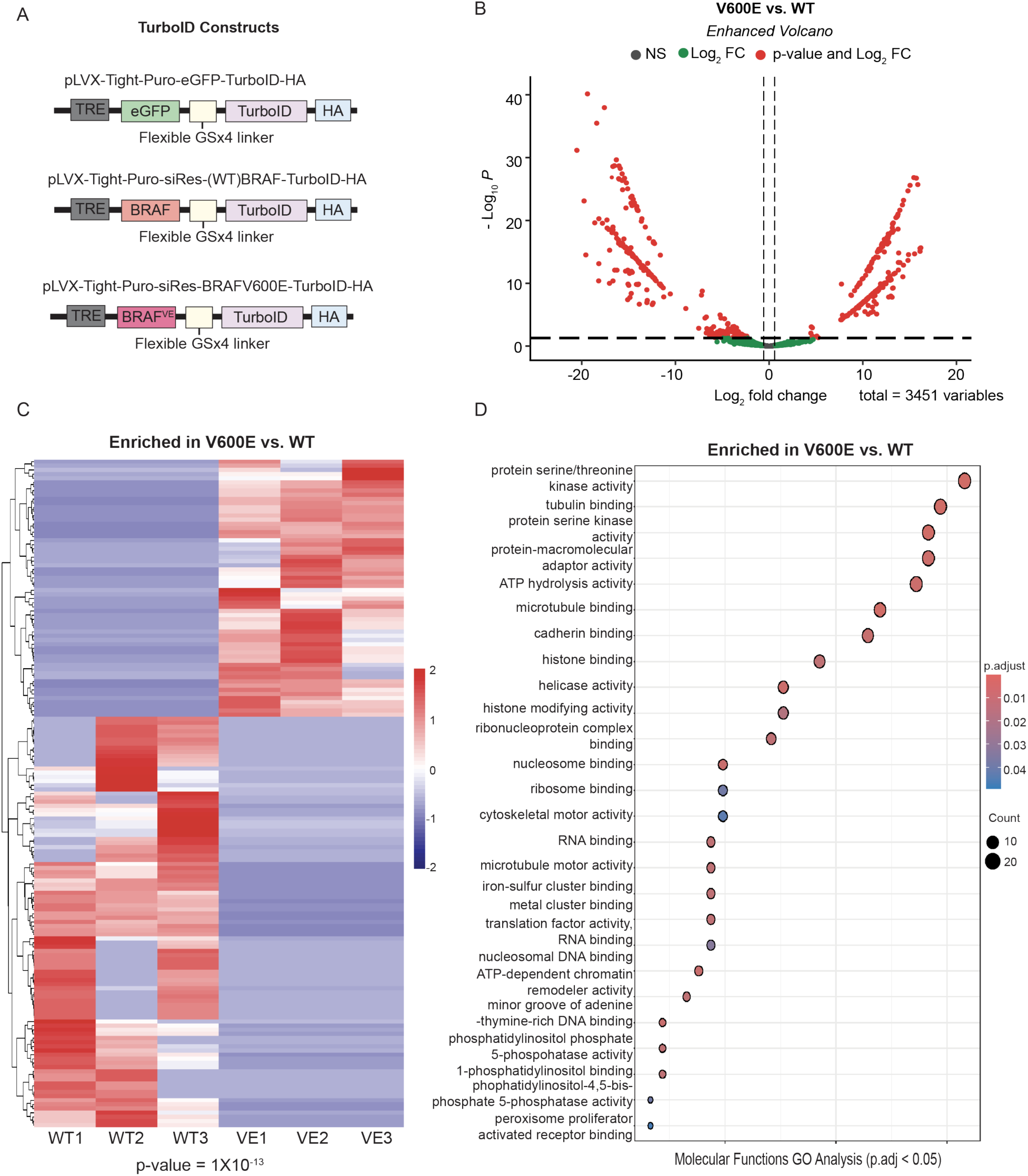
BRAF^V600E^ engages a unique set of protein binding partners. **A.** Design of pLVX-TurboID-HA vectors expressing eGFP, wild-type BRAF (WT), or BRAF^V600E^ (BRAF VE). TurboID-fusion constructs are expressed under the control of a tetracycline-response element (TRE) promoter. There is a flexible linker to allow for a larger radius of biotinylation and an HA tag for detecting construct expression. **B.** Volcano plot of differentially biotinylated proteins between BRAF wild-type and V600E. Red dots indicate proteins with padj < 0.05 and log2FC >2. **C.** Heatmap depicting BRAF interactors identified in BRAF-TurboID screen that are enriched in V600E vs. wild-type. Samples are in biological triplicates with the following conditions: wild-type BRAF-TurboID (WT) and BRAF^V600E^-TurboID (VE). **D.** Molecular functions gene ontology (GO) analysis comparing pathways enriched in BRAF^V600E^- TurboID hits compared to wild-type BRAF-TurboID hits. Pathways with adjusted p value<0.05 are depicted.

To identify BRAF protein-protein interactions with both normal and BRAF^V600E^, cells were treated with doxycycline to induce the expression of the siRNA-resistant TurboID constructs (Figure 1A) and then treated with siRNAs to endogenous BRAF. A doxycycline titration was performed to achieve equal expression of BRAF constructs. After culturing the cells for 24 hours in biotin-depleted media, cell culture media was supplemented with 50μM biotin for one hour, after which cells were harvested, lysed, and biotinylated proteins were affinity-captured on streptavidin-conjugated beads and subjected to tandem mass spectrometry in biological triplicates. We captured numerous proteins reported to interact with BRAF, including well-known interactors such as 14-3-3 and MEK1/2^43–48^. Significant protein interactors with BRAF^V600E^-TurboID (p < 0.05, Student’s t test) with at least a 2-fold enrichment relative to normal BRAF were prioritized for further analysis. This led to the identification of 228 proteins considered to interact more with BRAF^V600E^ compared to normal BRAF (padj < 0.05; log2FC > 1, Figure 1B). Interactors unique to both wild-type BRAF and BRAF^V600E^ were analyzed in triplicate (Figure 1C). Gene ontology (GO) analysis in molecular functions of BRAF^V600E^-specific interacting proteins revealed enrichment of expected pathways such as protein serine/threonine kinase activity, as well as pathways including protein-macromolecule adaptor activity and ATP hydrolysis activity (Figure 1D). Surprisingly, among hits enriched in the BRAF^V600E^-specific TurboID interactome was the tumor suppressor TP53 protein.

### BRAF-TurboID reveals novel interaction between BRAF and TP53

To further test the putative interaction of BRAF with TP53, we first repeated the streptavidin pulldowns in wild-type BRAF-TurboID and BRAF^V600E^-TurboID cell lines and confirmed that TP53 was enriched in the BRAF^V600E^-TurboID sample (Figure 2A). Moreover, Duolink in-situ proximity ligation assay (PLA) demonstrates that endogenous BRAF and TP53 interact in SKMEL-239 melanoma cells, which are heterozygous for the *BRAF^T1799A^*mutation. BRAF and pan 14-3-3 antibodies were used as a positive control (Figure 2B). For all experiments, stringent negative controls were performed, including: 1. Replicates without either primary antibody; 2. Replicates without secondary antibody probes; 3. Replicates with TP53 null cells. A375 cells, a human melanoma cell line that are homozygous for the *BRAF^T1799A^* mutation, display a strong BRAF<=>TP53 PLA signal (Figure 2C). To test if BRAF and TP53 interact in a non-BRAF^V600E^ melanoma cell line, we utilized Mel-9 cells, a human melanoma cell line expressing NRAS^Q61R^. In these cells, we observed a BRAF and TP53 interaction, albeit there was remarkably less BRAF<=>TP53 PLA signal in these cells (Figure 2D). Comparisons in different cancer cell lines that have different genetic alterations is challenging to control. Thus, in order to compare and quantify the BRAF<=>TP53 interaction in genetically matched cell lines, we utilized human epidermal melanocytes engineered to express a doxycycline-inducible BRAF^V600E^ oncoprotein kinase^49^. Upon doxycycline-induced expression of BRAF^V600E^ in primary epidermal melanocytes, we observed a significant increase in the PLA signal between BRAF^V600E^ and TP53 (Figures 2E and 2F). This induction of BRAF<=>TP53 association indicates a previously unrecognized spatial proximity between these two proteins in melanocytes and melanoma cells.

**Figure 2:**
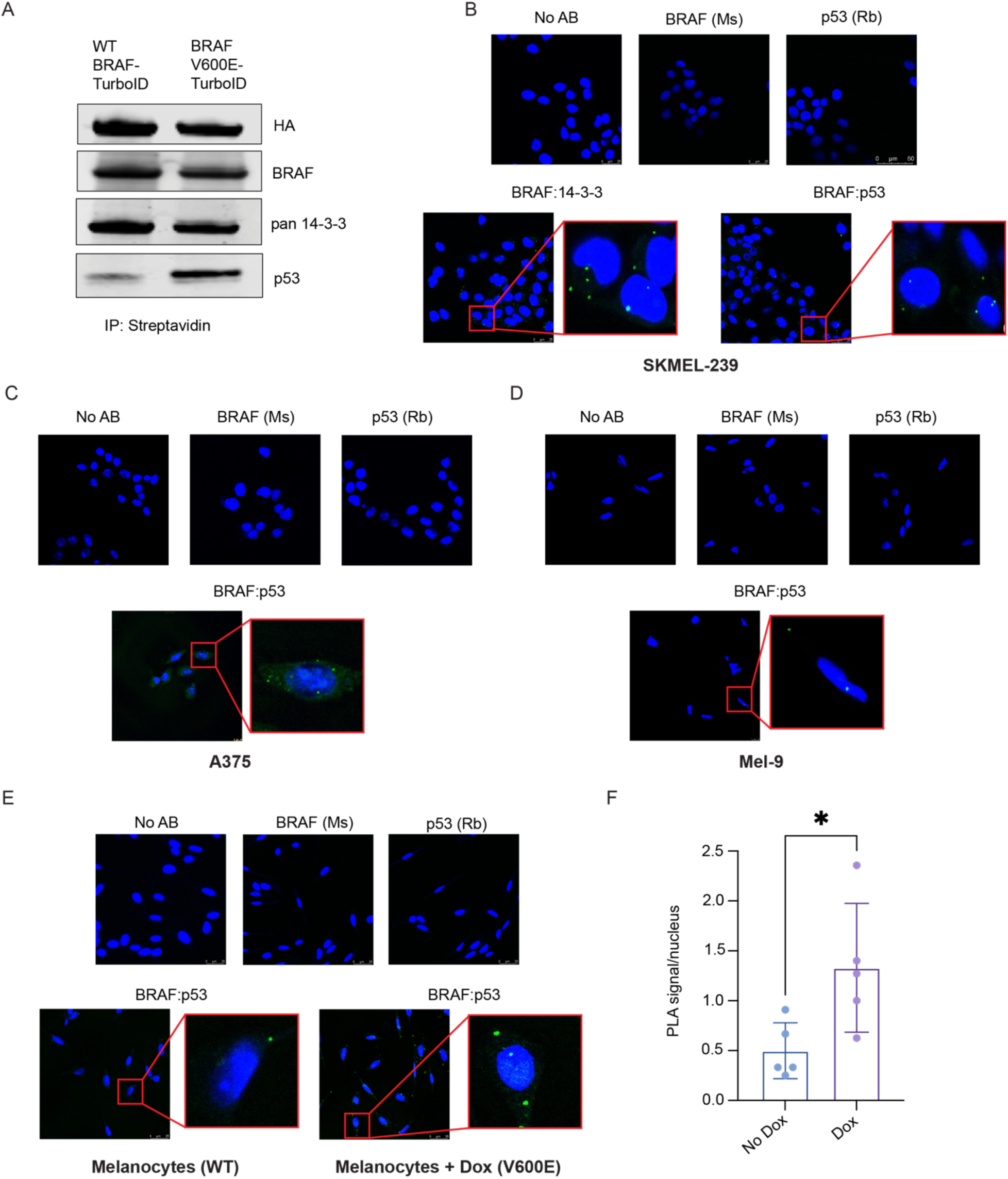
BRAF^V600E^ exhibits enhanced BRAF and TP53 interaction. **A.** Immunoblots demonstrating TP53 pulldown with BRAF through streptavidin pulldown from BRAF-TurboID (wild-type and V600E) whole cell lysates. **B.** Proximity ligation assays (PLA) in SKMEL-239 human melanoma cell line stained with DAPI (blue) showing no antibody and single antibody controls and cytoplasmic PLA signal (green) for BRAF:14-3-3 positive control and BRAF:TP53 (Scale bar = 25μM. Representative images shown from n=3 biological replicates). **C.** PLA in human melanoma cell line A375 stained with DAPI (blue) showing no antibody and single antibody controls and cytoplasmic PLA signal (green) for BRAF:TP53 (Scale bar = 25μM. Representative images shown from n=3 biological replicates). **D.** PLA in human melanoma cell line Mel-9 (NRAS^Q61R^) stained with DAPI (blue) showing no antibody and single antibody controls and cytoplasmic PLA signal (green) for BRAF:TP53 (Scale bar = 25μM. Representative images shown from n=3 biological replicates). **E.** PLA in human epidermal melanocytes with a doxycycline-inducible BRAF^V600E^ stained with DAPI (blue) showing no antibody and single antibody controls and cytoplasmic PLA signal (green) for BRAF:TP53 (Scale bar = 25μM. Representative images shown from n=3 biological replicates). **F.** Quantification of BRAF:TP53 PLA signal from primary epidermal melanocytes with and without the addition of doxycycline to turn on the expression of BRAF^V600E^, reported as average PLA signals per nucleus (quantification of n=4-5 fields of view from each of n=3 biological replicates; unpaired t-test p=0.0295).

### TP53 interacts with BRAF through its DNA-binding domain

To determine the amino acid residues and the interaction interface between BRAF and TP53, we engineered a series of TP53 deletion mutants (Figure 3A) with FLAG-tagged TP53 and GST-tagged normal BRAF and BRAF^V600E^ ^15^. TP53 is comprised of two N-terminal transactivation domains (TAD) consisting of residues 1-61; a proline-rich domain (PRD) of residues 64-92; a DNA-binding domain (DBD) of residues 96-262, which is linked to a tetramerization domain (TET) from residues 324-356 through a nuclear localization sequence (NLS); a tetramerization domain (TET); a nuclear export sequence (NES); and lastly, a C-terminal regulatory domain (CTD) which consists of residues 364-393^50^. Co-immunoprecipitation experiments with a FLAG antibody reveal that the DNA-binding domain of TP53 is required for the interaction with both normal BRAF and BRAF^V600E^ (Figures 3B-C). Reciprocal pulldowns of GST-BRAF confirmed this requirement of the TP53 DNA-binding domain (data not shown). We then expressed FLAG-TP53 expressing no DNA-binding domain or the DNA-binding domain alone (Figure 3D). Indeed, BRAF co-immunoprecipitated with the DNA-binding domain alone but not with the full-length TP53 construct missing the DNA-binding domain (Figure 3E), confirming the requirement of the DNA-binding domain of TP53 for interactions with BRAF.

**Figure 3:**
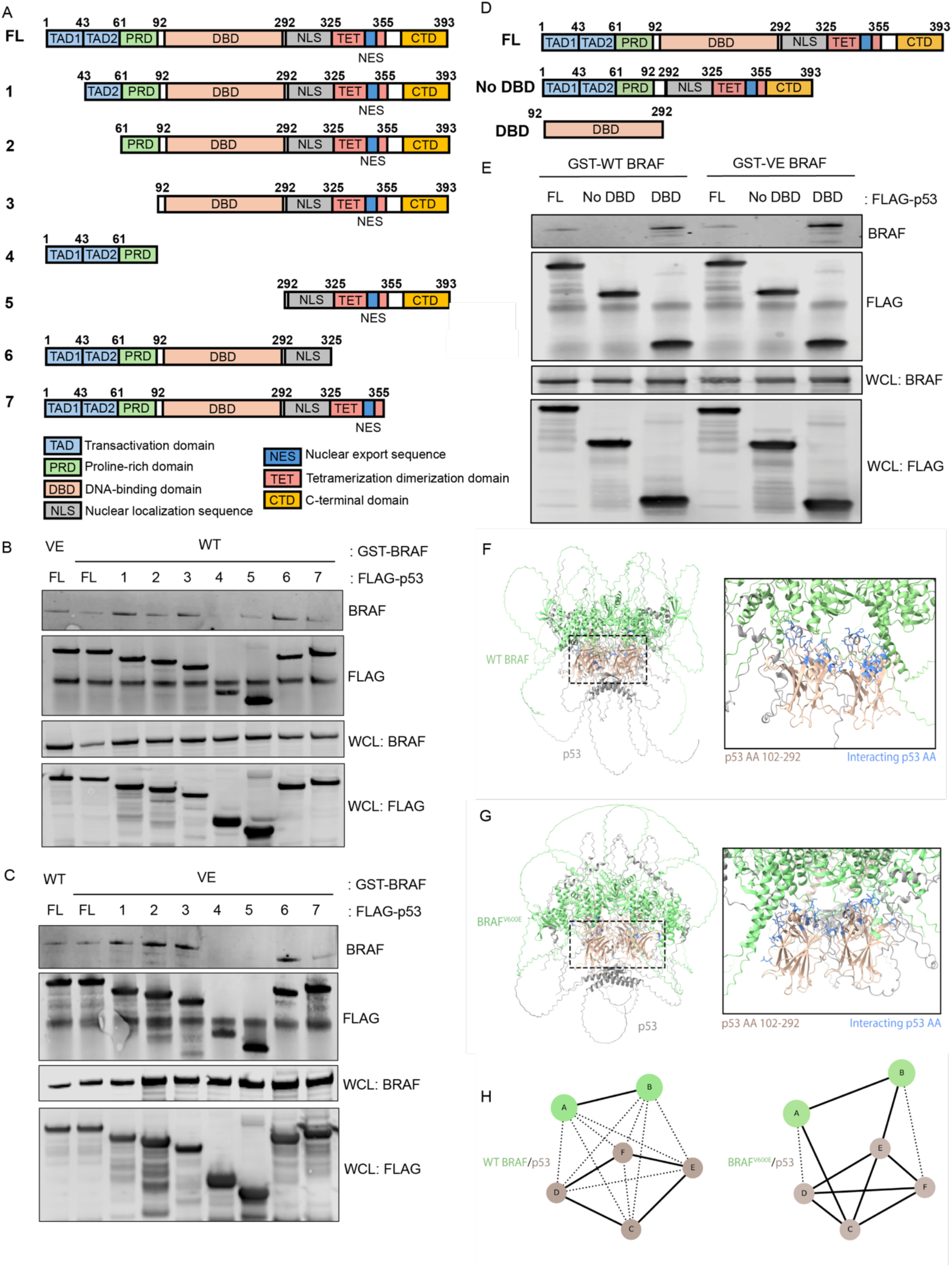
TP53 interacts with BRAF through its DNA binding domain. **A.** Schematic of FLAG-tagged TP53 deletion mutants. Mutants are labeled in numerical order in which they are depicted in the pulldowns (FL = full length TP53). **B.** Immunoblots depicting FLAG immunoprecipitations of TP53 deletion mutants co-expressed with GST-wild-type BRAF constructs (WT). WCL = whole cell lysate. **C.** Immunoblots depicting FLAG immunoprecipitations of TP53 deletion mutants co-expressed with GST-BRAF^V600E^ constructs (VE). WCL = whole cell lysate. **D.** Schematic of FLAG-tagged TP53 mutants with DBD alterations. **E.** Immunoblots depicting FLAG immunoprecipitations of TP53 DBD mutants co-expressed with GST-wild-type BRAF or GST-BRAF^V600E^ constructs. WCL = whole cell lysate. **F.** AlphaFold 3 prediction of wild-type BRAF (green) and TP53 (tan), interactions with interacting domains within the TP53 DNA-binding domain highlighted in blue. ipTM score of 0.29 and PTM score of 0.34. **G.** AlphaFold 3 prediction of BRAF^V600E^ (green) and TP53 (tan) interaction with interacting domains within the TP53 DNA-binding domain highlighted in blue. ipTM score of 0.29 and PTM score of 0.35. **H.** AlphaFold 3-based models of wild-type BRAF-TP53 complex (left) or BRAF^V600E^-TP53 complex (right) were analyzed for interaction interfaces in ChimeraX. Dark lines represent chain to chain interfaces of greater surface area. The dotted lines represent interfaces that are smaller than half of the largest interface. Chains A and B (green) represent BRAF sequences and chains C, D, E, and F represent TP53 surfaces.

To complement our *in vitro* data with the TP53 truncation mutants, we employed AlphaFold3 to model the BRAF^V600E^ and TP53 interaction. This approach corroborated our pull-down data in Figures 3B, C, and E by predicting the DNA binding domain of TP53 as the interacting surface with the BRAF^V600E^ kinase domain (Figures 3F-G). Furthermore, a comparison of the normal BRAF- and BRAF^V600E^-TP53 complexes suggests that structural changes caused by the V600E substitution are predicted to expand the interaction surface with 26 interacting residues on TP53 and 22 on normal BRAF versus 91 interacting residues on TP53 and 31 on BRAF^V600E^ (Figure 3G). Figure 3H depicts these changes in an interface diagram (ChimeraX) in which the darker lines represent side chain-to-side chain interfaces of greater surface area between TP53 and BRAF. Thus, the structure prediction validated our pulldown data (Figure 3B, C, E), suggesting that V600E substitution-induced conformational changes enhance the BRAF^V600E^-TP53 interaction.

### BRAF and TP53 colocalize in melanoma cells and epidermal melanocytes driven by oncogenic BRAF

TP53 is known for its canonical transcription factor functions upon DNA binding in the nucleus. However, TP53 is reported to shuttle to the cytoplasm through the nuclear pore complex^51^ and has also been suggested to localize at the outer mitochondrial membrane^52,53^. However, the signals for TP53 to translocate outside of the nucleus are not fully elucidated. Thus, we aimed to understand how TP53 localization changes with BRAF activation and evaluate if TP53 colocalizes with BRAF in the cell. To visualize the interaction between BRAF and TP53, we performed immunofluorescence microscopy with human melanoma cell lines driven by either BRAF^V600E^ or NRAS^Q61R^, as well as HEK293 cells. Interestingly, in cell lines not expressing BRAF^V600E^ (HEK293 and Mel-9), TP53 was primarily localized in the nucleus (Figures 4A and 4B, respectively). By contrast, BRAF^V600E^-expressing cell lines (SKMEL-239 and A375), demonstrated TP53 localized throughout the nucleus and the cytoplasm and colocalized with BRAF (Figures 4C and 4D, respectively). Quantification of BRAF and TP53 or TP53 and DAPI colocalization revealed increased BRAF and TP53 colocalization in the melanoma cells driven by BRAF^V600E^ (Figure 4E) and inversely, less nuclear TP53 (Figure 4F). Specifically, the melanoma cells driven by BRAF^V600E^had ∼20% more BRAF and TP53 colocalization and ∼10-20% less nuclear TP53. In the primary human epidermal melanocytes described above, addition of doxycycline to express BRAF^V600E^ resulted in a reduction of nuclear TP53 staining in around half of the cells and increased TP53 within the cytoplasm (Figures 4G-J). Notably, some cells maintained nuclear TP53 staining, while other cells display a nearly a complete reduction of nuclear TP53 localization.

**Figure 4:**
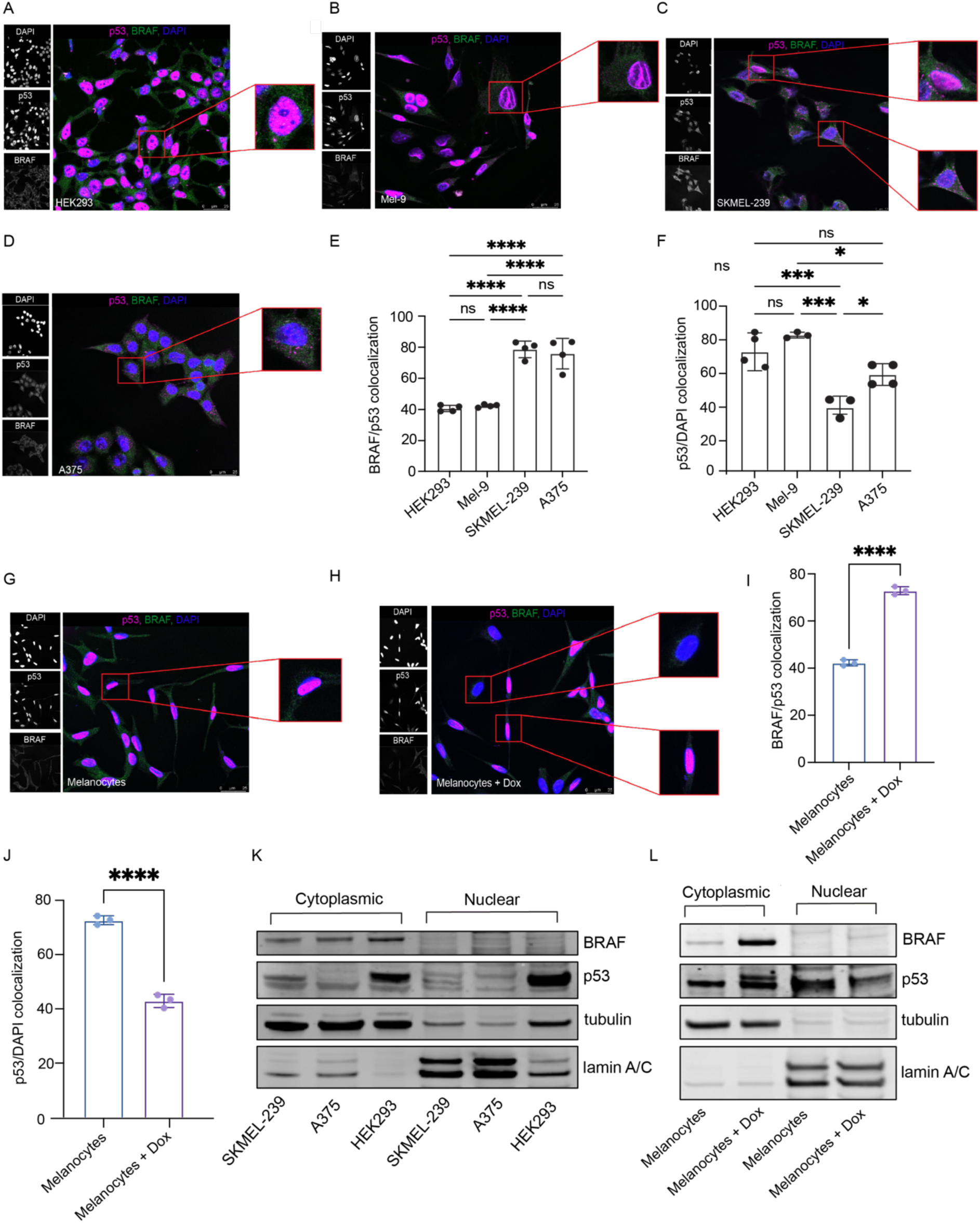
BRAF and TP53 colocalize in the cytoplasm Immunofluorescence microscopy depicting TP53 (magenta) and BRAF (green) localization in: **A.** HEK293 cells (wild-type BRAF). **B.** Mel-9 cells (NRAS^Q61R^, wild-type BRAF). **C.** SKMEL-239 cells (heterozygous for BRAF^V600E^). **D.** A375 cells (homozygous for BRAF^V600E^). **E.** Quantification of colocalization of BRAF and TP53 in the cell lines listed above (quantification of n=4-5 fields of view from each of n=3 biological replicates; one-way ANOVA with multiple comparisons. **** = adjusted p-valued <0.0001). **F.** Quantification of TP53 localization in the nucleus by DAPI staining in the cell lines listed above (quantification of n=4-5 fields of view from each of n=3 biological replicates; one-way ANOVA with multiple comparisons. * = adjusted p-valued <0.03, *** = adjusted p-value <0.0009). Immunofluorescence microscopy depicting TP53 (magenta) and BRAF (green) localization in: **G.** Primary epidermal melanocytes expressing wild-type BRAF with no doxycycline. **H.** Primary epidermal melanocytes expressing BRAF^V600E^ with the addition of doxycycline. **I.** Quantification of TP53 and BRAF colocalization in the epidermal melanocytes (quantification of n=4-5 fields of view from each of n=3 biological replicates; unpaired t-test p-value <0.0001). **J.** Quantification of TP53 localization in the nucleus by DAPI staining (quantification of n=4-5 fields of view from each of n=3 biological replicates; unpaired t-test p-value <0.0001). **K.** Immunoblots detecting BRAF and TP53 localization in the cytoplasm vs. nucleus after cell fractionation of SKMEL-239 cells, A375 cells, and HEK293 cells. **L.** Immunoblots detecting BRAF and TP53 localization in the cytoplasm vs. nucleus after cell fractionation of primary epidermal melanocytes with and without the addition of doxycycline to turn on BRAF^V600E^ expression.

To complement the immunofluorescence microscopy experiments, we performed cell fractionation to separate the nuclear and cytoplasmic components. In HEK293 cells, BRAF is predominantly cytoplasmic and TP53 is detected in both fractions, but more abundant in the nuclear fraction (Figure 4K). However, in the SKMEL-239 and A375 melanoma cells, driven by BRAF^V600E^, TP53 is largely detected in the cytoplasm and not the nuclear fraction (Figure 4K). Utilizing the primary epidermal melanocytes, induction of BRAF^V600E^ expression results in more cytoplasmic TP53 and less nuclear TP53, compared to normal BRAF (Figure 4L). Hence, these data suggest that BRAF^V600E^ expression leads to TP53 accumulation in the cytoplasm, therefore leading to less nuclear TP53.

### BRAF^V600E^ expression decreases TP53 activity

Melanoma has long been noted for a low frequency of *TP53* mutations, despite the fact that it is an aggressive disease. Numerous studies have shown that even though most melanomas retain normal *TP53*, the TP53 protein does not respond to appropriate biochemical or biological cues^32,34,35^. Thus, we sought to evaluate TP53 activity in our melanoma cell lines. First, we tested that as BRAF^V600E^ expression increases, the BRAF<=>TP53 interaction increases concurrently (Figures 5A-B). However, as BRAF^V600E^ expression rises, so does total TP53 protein expression (Figure 5C). This stabilization of wild-type TP53 is typically induced by signals that activate TP53^54–56^. We observe elevated TP53 expression in our BRAF wild-type HEK293 cells when treating with the MDM2 inhibitor, Nutlin-3a (Figure 5D). However, we noted that TP53 is unable to be activated or stabilized by treatment with Nutlin-3a in our BRAF^V600E^ melanoma cell lines (Figures 5E-F). Consistently, Nutlin-3a increased TP53 expression in the primary human epidermal melanocytes expressing normal BRAF, but not with melanocytes expressing BRAF^V600E^ (Figures 5G-H). We also subjected the epidermal melanocytes with or without doxycycline treatment to bulk RNA-sequencing (RNA-seq). Importantly, expression of *TP53* and *MDM2* mRNAs remained similar in both cell types (Figure 5I).

**Figure 5:**
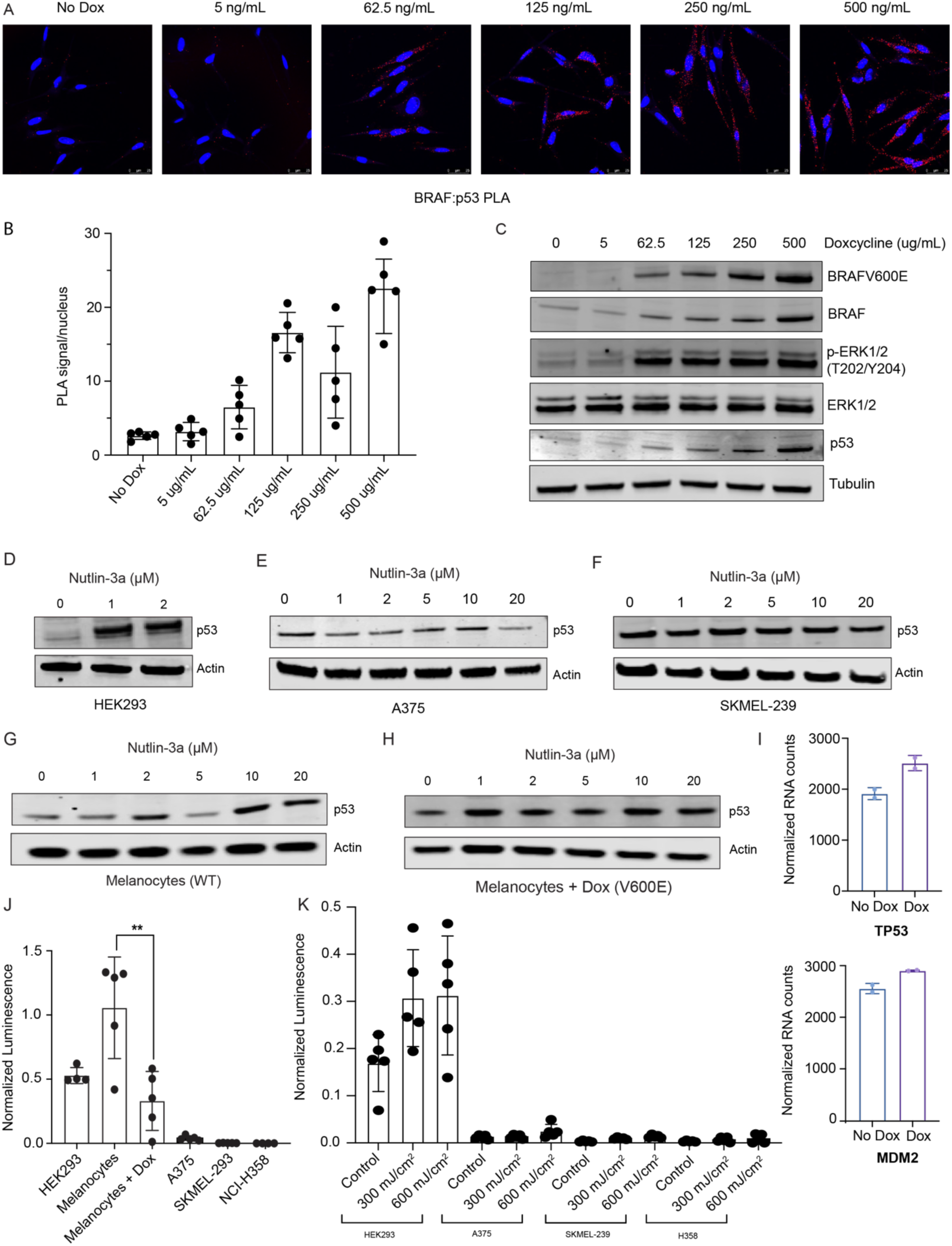
BRAF^V600E^ limits TP53 activation. **A.** PLA in human epidermal melanocytes with a doxycycline-inducible BRAF^V600E^ with increasing doxycycline concentrations, stained with DAPI (blue) and cytoplasmic PLA signal (red) for BRAF:TP53 (Scale bar = 25μM. Representative images shown from n=3 biological replicates). **B.** Quantification of BRAF:TP53 PLA signal from primary epidermal melanocytes with and without increasing doses of doxycycline to induce BRAF^V600E^ expression, reported as average PLA signals per nucleus (quantification of n=4-5 fields of view from each of n=3 biological replicates). **C.** Immunoblot analyses of increasing BRAF^V600E^ expression and increased phosphorylated ERK with increasing doxycycline. **D.** Immunoblots of HEK293 cells treated with increasing doses of Nutlin-3a. **E.** Immunoblots of A375 cells treated with increasing doses of Nutlin-3a. **F.** Immunoblots of SKMEL-239 cells treated with increasing doses of Nutlin-3a. **G.** Immunoblots of primary epidermal melanocytes expressing wild-type BRAF treated with increasing doses of Nutlin-3a. **H.** Immunoblots of primary epidermal melanocytes with doxycycline expressing BRAF^V600E^ treated with increasing doses of Nutlin-3a. **I.** Normalized RNA counts of *TP53* and *MDM2* from RNA-sequencing of epidermal melanocytes with doxycycline-inducible BRAF^V600E^. **J.** Relative TP53 activity in various human cell lines. The PG13-luc luciferase reporter for TP53 activity was co-transfected with a renilla control vector and dual luciferase activities were assessed. (Individual dots represent five technical replicates. One-way ANOVA for multiple comparisons, ** = adjusted P value = 0.0003). **K.** Relative TP53 activity in human cell lines exposed to increasing doses of UVB as indicated. The PG13-luc luciferase reporter for TP53 activity was co-transfected with a renilla control vector and dual luciferase activities were assessed. (Individual dots represent five technical replicates).

Next, a TP53 responsive luciferase reporter system revealed extremely low TP53 activity in cell lines with BRAF^V600E^ expression. This low TP53 activity in our BRAF-driven melanoma cell lines was comparable to our negative control cells, NCI-H358, which are human non-small cell lung cancer cells that have a *TP53* deletion. These cells have no TP53 reporter activity and serve as our baseline. In the melanocytes with expression of normal BRAF, they have relatively higher levels of TP53 activity at baseline, which is significantly decreased following doxycycline treatment to induce BRAF^V600E^ expression (Figure 5J). Importantly, the expression of BRAF^V600E^ leads to ∼50% less TP53 transcriptional activity compared to melanocytes expressing normal BRAF.

UVB radiation is well-established to activate TP53 and is a strong inducer of apoptosis in keratinocytic and melanocytic cells^57^. Therefore, we evaluated TP53 activity in our cell lines after high-dose UVB radiation (300-600mJ/cm^2^). Here, we demonstrate induction of TP53 activity in a dose-dependent manner in HEK293 cells, but UVB radiation fails to activate TP53 in our human melanoma cells driven by BRAF^V600E^ (Figure 5K). Thus, although TP53 is expressed in melanoma cell lines driven by BRAF^V600E^, TP53 is unable to be activated through conventional techniques such as MDM2 inhibition or UVB radiation.

### TP53 mutations are not necessary in BRAF-driven melanoma

We propose that BRAF^V600E^ expression prevents TP53 from translocating to the nucleus and thus prevents normal TP53 function in melanoma cells (Figure 6A). *TP53* mutations only occur in ∼17% of cutaneous melanomas, a much lower frequency compared to non-melanoma skin cancers where the more than half of patients exhibit TP53 alterations^58,59^. Interestingly, mutations or loss of the *TP53* gene in cutaneous melanoma cases do not change the probability of survival for these patients (Figure 6B). This is surprising given that alterations in *TP53* are known to accelerate lung, thyroid, breast and other cancers^60–64^. Furthermore, *TP53* loss or mutations are mutually exclusive with BRAF^V600E^ mutations in the cancers commonly driven by oncogenic BRAF (cutaneous melanoma, lung adenocarcinoma, colorectal carcinoma, and thyroid cancer) (Figure 6C). Thus, the interaction between TP53 and oncogenic BRAF may stabilize TP53 and prevents it from translocating back to the nucleus to elicit its tumor suppressor function.

**Figure 6:**
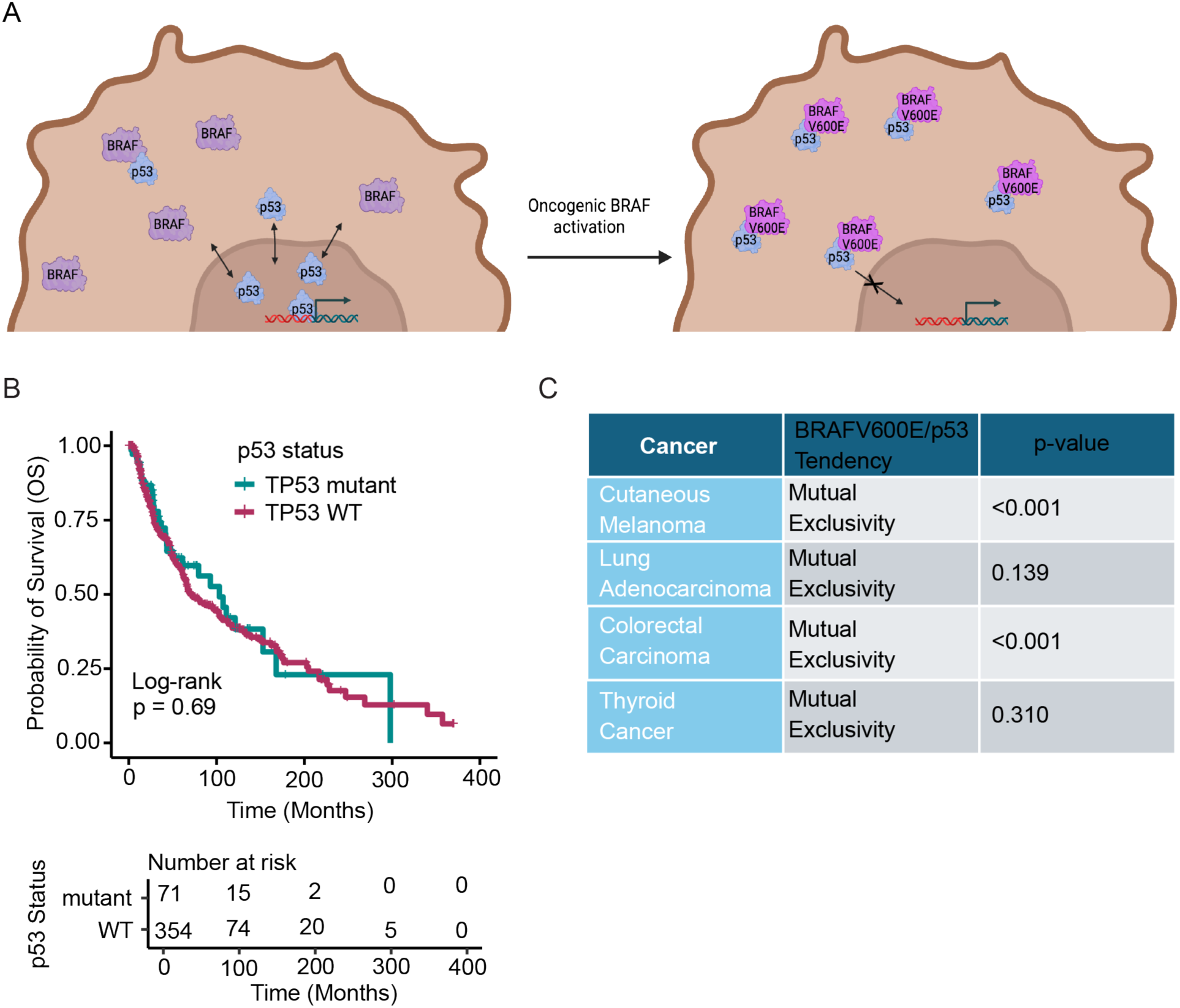
TP53 mutations are not necessary in BRAF-driven cancers. **A.** Proposed model of BRAF^V600E^ acting as a molecular “trap” and preventing TP53 from translocating to the nucleus. **B.** Kaplan-Meier curve of PanCancer TCGA skin cutaneous melanoma samples stratified by presence or absence of a *TP53* mutation. Log-Rank test. **C.** Mutual exclusivity chart of cancers commonly driven by BRAF with tendency of mutual exclusivity or co-occurrence with BRAF^V600E^ and TP53 mutation or loss with p-value from one-sided Fisher’s Exact Test (data from cBioPortal, PanCancer TCGA).

## DISCUSSION

Tightly regulated signaling within the RAS>RAF>MEK>ERK pathway is critical for maintaining normal cellular functions and survival. Importantly, it is most frequently mutated cell signaling pathway in cancer. While extremely well studied, many questions remain regarding oncogenic BRAF functions compared to normal. Moreover, prior attempts to investigate the interactomes of the RAF family kinases^14,17,18^ using conventional techniques often fail to capture weak or transient interactions. Resistance to BRAF^V600E^ targeted therapies remains a major therapeutic obstacle, limiting both the depth and durability of responses in melanoma patients. Identifying proteins that interact with BRAF^V600E^ can provide critical insights into its cellular roles driving malignancy. Thus, we aimed to better understand how oncogenic BRAF functions with the ultimate goal of identifying BRAF^V600E^ interactors that are critical to its oncogenic activity. Intriguingly, while there was overlap in protein-protein interactions, wild-type BRAF and BRAF^V600E^ also exhibited many different protein-protein interactions.

*TP53* is the most frequently mutated gene in cancer. TP53 functions as the “guardian of the genome”, activating numerous cellular processes including apoptosis, cell cycle arrest, and DNA repair among others^25,54,65^. This cell protective activity makes TP53 an ideal target for inactivation by cancer cells. Indeed, alterations in TP53 occur in over 50% of all cancer cases^20–22^. Moreover, alterations in *TP53* accelerate lung, thyroid, breast and other cancers^60^. Intriguingly, *TP53* is much less frequently lost or mutated in melanoma, as the majority of melanomas harbor wild-type *TP53* (>80%). Moreover, several studies have shown that when melanomas express wild-type *TP53*, its tumor suppressor activity is inhibited^31,32,34–36,66,67^.

Our data reveal important changes in TP53 function as a consequence of BRAF^V600E^ expression. As BRAF^V600E^ is expressed, the BRAF-TP53 interaction increases and TP53 appears to translocate to the cytoplasm. Additionally, TP53 is unable to be activated in response to MDM2 inhibition as well as through UVB radiation, and TP53 activity is almost undetectable in BRAF^V600E^ melanoma cell lines. Importantly, TP53 activity decreases with BRAF^V600E^ expression alone. Thus, we hypothesize that BRAF^V600E^ interacts with TP53 in the cytoplasm and essentially acts as a “trap”, preventing TP53 from eliciting its nuclear functions in order to promote melanoma cell survival.

Based on the results from our TurboID screen, BRAF can interact with many proteins, and we have yet to elucidate the full mechanism by which BRAF may interact with TP53. Other proteins identified in our TurboID screen could facilitate this interaction including AHNAK (desmoyokin), TP53BP2 (ASPP2), and other TP53 binding partners. Importantly, both TP53 and BRAF interact with the chaperone protein HSP90^68,69^. We were unsuccessful in knocking down and inhibiting HSP90 as it has numerous critical functions in cells. However, it’s possible that HSP90 could be brokering this interaction. Additionally, research in melanoma has shown that MDM4 (MDMX) plays a role in regulating TP53^38^. We did not detect MDM4 as a hit in our TurboID screen and the mRNA expression levels of MDM4 were the same in the melanocytes with and without expression of BRAF^V600E^. However, there are likely numerous cellular mechanisms that are facilitating the suppression of TP53 activity in melanoma cells. Finally, our results validate dysregulated TP53 function in melanoma cells driven by oncogenic BRAF and characterize a novel interaction between BRAF and TP53, which favors BRAF^V600E^. Importantly, we show that BRAF^V600E^ expression alone alters TP53 localization and decreases TP53 transactivation activity. Our identification of a functional BRAF-TP53 interaction provides new mechanistic insight into the disruption of the tumor suppressive function of TP53 in melanoma and other BRAF-driven cancers.

## MATERIALS AND METHODS

### Cell Culture

All cell lines were routinely tested for mycoplasma contamination. All human lung cancer cell lines were cultured in RPMI (Roswell Park Memorial Institute) 1640 (Gibco 11875-093) supplemented with 10% fetal bovine serum (Gibco 10438-026) and 1% penicillin plus streptomycin (Gibco 15140-122). Melanoma cells were cultured in DMEM/F12 (Dulbecco’s Modified Eagle Medium F12) (Gibco 11330-032) with 10% fetal bovine serum and 1% penicillin plus streptavidin. Primary epidermal melanocytes were cultured in Melanocyte medium (Thermo Fisher Scientific M254500) with HMGS (Thermo Fisher Scientific S0025).

### Lentiviral Transduction

HEK293T cells were seeded 24 hours before transduction in DMEM/F12 (10% FBS 1% p/s) and Lipofectamine 3000 kit (Invitrogen L3000015) was used for all lentiviral generation. All virus-containing supernatants were filtered through 0.45 micro filters before use. To increase the efficiency of infection, 10ug/mL of polybrene (MilliporeSigma TR-1003-G) was supplemented in the virus-containing media when added to cells. Cells were selected for successful infection through antibiotic selection with the corresponding antibiotics (puromycin and blasticidin).

### Cloning and Site Directed Mutagenesis

See table 1 for a list of genetic TurboID constructs used in this study.

**Table 1.**

| Reagent type | Designation | Source or<br>Reference | Identifiers | Additional<br>Information |
| --- | --- | --- | --- | --- |
| Recombinant<br>DNA reagent | pLVX-Tight-Puro<br>backbone | PMID: 37772788 |  | Sundquist Lab |
| Recombinant<br>DNA reagent | pLVX-eGFP-TurboID-HA |  |  | Sundquist Lab |
| Recombinant<br>DNA reagent | pLVX-BRAF-TurboID-HA |  |  |  |
| Recombinant<br>DNA reagent | pLVX-BRAFV600E-TurboID-HA |  |  |  |
| Recombinant<br>DNA reagent | RCAS-BRAF | PMID: 19855433 |  | Holmen Lab |
| Recombinant<br>DNA reagent | GST-BRAF (WT) | PMID: 35512704 |  | Fu Lab |
| Recombinant<br>DNA reagent | GST-BRAF<br>(V600E) | PMID: 35512704 |  | Fu Lab |
| Recombinant<br>DNA reagent | FLAG-TP53 | PMID: 35512704 |  | Fu Lab |
| Recombinant<br>DNA reagent | PG13-luc(wt<br>TP53 binding<br>sites) | Addgene | Plasmid # 16442 |  |
| Recombinant<br>DNA reagent | pRL-CMV-Renilla control<br>vector | Promega | Cat# E2261 |  |
| Recombinant<br>DNA reagent | pTRIPZ-BRAFV600E |  |  | Judson-Torres<br>Lab |

To generate the BRAF-TurboID constructs (pLVX-siRes-BRAF-TurboID-HA), PCR fragments from RCAS-BRAF (gift from Dr. Sheri Holmen) and pLVX-TurboID (gift from Dr. Wes Sundquist) were assembled by overlapping ends using Gibson assembly master mix (New England Biolabs (NEB). The pLVX-Tight-Puro construct was linearized through a restriction enzyme double digestion with BamHI and MluI. BRAF (wild-type and V600E) were amplified with primers (5’- agatcgcctggccaccatggcggcgctg-3’) and (5’-cctgatcctggacaggaaacgcaccatatcc-3’), while TurboID was amplified with primers (5’-cctgtccaggatcaggaagcggatcagga-3’) and (5’- cctacccggtagaattcaCTAAGCGTAGTC-3’). Linearized and fragment DNA were analyzed using 1% (w/v) agarose gels. After Gibson assembly, One Shot Stbl3 chemically competent cells (Thermo Fisher Scientific C737303) were used for transformation. Qiagen maxi prep kits (12162) were used to harvest DNA according to the manufacturers protocol. Sanger sequencing was performed on the purified plasmids by the HSC DNA Sequencing Core at the University of Utah.

The RCAS-BRAF plasmids (WT and V600E) were engineered to be siRNA resistant with 6 silent point mutations (5’-gtggcatggtgatgtggca-3’ to 5’-AtggcaCggGgaCgtAgcT-3’). These mutations were engineered to confer resistance to siRNA’s targeting endogenous BRAF 5’- AAGUGGCAUGGUGAUGUGGCA-3’^70^. The mutations were introduced at the same time by site-directed mutagenesis using nonoverlapping primers with NEB Q5 SDM protocol and were designed by the NEBaseChanger program (5’-ggacgtagctGTGAAAATGTTGAATGTGAC-3’ and 5’-ccgtgccatttTCCCTTGTAGACTGTTCC-3’). The New England Biolabs Q5 Site-Directed Mutagenesis Kit (E0554S) was utilized to create the DNA changes and DNA was transformed using their NEB 5-alpha Competent *E. coli* cells. Bacterial DNA was harvested using Qiagen mini prep kits (27104). All site-directed alterations were confirmed by DNA sequencing. GST-BRAF (WT and V600E) plasmids were a gift from Dr. Haian Fu.

### Immunoblotting

Cells were washed twice with ice-cold PBS and scraped in 1mL of ice-cold PBS. Cells were pelleted by centrifugation for 10 seconds at 13,000 rpm at 4°C. Supernatant was removed, and cell pellets were resuspended in 25-200μL (depending on pellet size) in RIPA buffer (50mM Tris pH 8.0, 150mM NaCl, 0.5mM EDTA, 10mM NAF, 0.1% SDS, 0.5% sodium deoxycholate, 1% NP-40 substitute) plus HALT protease inhibitor cocktail (Thermo Fisher Scientific 78430) at 2X concentration and incubated on ice for 20 minutes. Cellular debris was pelleted for 10 minutes at 4°C and protein concentration was quantified using a bicinchoninic acid assay (BCA) (Thermo Fisher Scientific 23250). 25μg of samples were equally loaded for SDS-PAGE and transferred on nitrocellulose membranes. Membranes were blocked for one hour in 5% BSA and probed with primary antibodies (Table 2) overnight in a cold room. Secondary antibodies (Table 2) used were diluted and incubated at RT for two hours. Membranes were imaged and analyzed on an Odyssey® CLx Infrared Imaging System (LI-COR®) and Image Studio Software (LI-COR®).

**Table 2.**

| Reagent type | Designation | Source or Reference | Identifiers | Additional Information |
| --- | --- | --- | --- | --- |
| Antibody | Anti-β-actin<br>(mouse monoclonal) | Cell Signaling Technology | Cat# 3700 | WB (1:10,000) |
| Antibody | Anti-BRAF F-7<br>(mouse monoclonal) | Santa Cruz Biotechnology | Cat# sc-5284 | WB (1:500) |
| Antibody | Anti-BRAF<sup>V600E</sup><br>(mouse monoclonal) | Abcam | Cat# ab228461 | WB (1:1,000) |
| Antibody | Anti-Cyclin D1<br>(rabbit polyclonal) | Cell Signaling Technology | Cat# 2922 | WB (1:1,000) |
| Antibody | Anti-DYKDDDDK Tag<br>(rabbit monoclonal) | Cell Signaling Technology | Cat# 14793 | WB (1:1,000) |
| Antibody | Anti-p44/42 ERK1/2<br>(mouse monoclonal) | Cell Signaling Technology | Cat# 4696 | WB (1:1,000) |
| Antibody | Anti-phospho-p44/42 | Cell Signaling Technology | Cat# 4377 | WB (1:1,000) |
|  | Thr202/Tyr204<br>ERK1/2 (rabbit<br>monoclonal) |  |  |  |
| Antibody | Anti-GAPDH<br>(mouse<br>monoclonal) | Cell Signaling<br>Technology | Cat# 97166 | WB (1:1,000) |
| Antibody | Anti-HA (mouse<br>monoclonal) | Cell Signaling<br>Technology | Cat# 2367 | WB (1:1,000) |
| Antibody | Anti-HSP90<br>(rabbit<br>polyclonal) | Cell Signaling<br>Technology | Cat# 4875 | WB (1:1,000) |
| Antibody | Anti-lamin A/C<br>(mouse<br>monoclonal) | Cell Signaling<br>Technology | Cat# 4777 | WB (1:1,000) |
| Antibody | Anti-pan 14-3-3<br>(rabbit<br>polyclonal) | Cell Signaling<br>Technology | Cat# 8312 | WB (1:1,000) |
| Antibody | Anti-TP53<br>(mouse<br>monoclonal) | Cell Signaling<br>Technology | Cat# 2524 | WB (1:1,000) |
| Antibody | Anti-TP53 (rabbit<br>monoclonal) | Cell Signaling<br>Technology | Cat# 2527 | WB (1:1,000) |
| Antibody | Anti-β-Tubulin<br>(rabbit<br>polyclonal) | Cell Signaling<br>Technology | Cat# 2146 | WB (1:5,000) |
| Antibody | IRDye 800CW<br>Goat anti-Rabbit<br>IgG | LI-COR | Cat# 926-32211 | WB (1:20,000) |
| Antibody | IRDye 680LT<br>Donkey anti-Mouse IgG | LI-COR | Cat# 926-68022 | WB (1:20,000) |
| Fluorsecent<br>streptavidin<br>conjugate | IRDye 800CW<br>Streptavidin | LI-COR | Cat# 926-32230 | WB (1:1,000) |

### TurboID

HEK293 TetOn cells were purchased from Takara (631182). Stable TurboID cell lines were generated in the HEK293 TetOn cells with lentiviral transduction. After a week of selection in puromycin (10μg/mL), cells were plated in single cell suspensions and clones were matched among conditions based on TurboID construct expression levels. For the screen, TurboID expressing cell lines were seeded in triplicate onto 15cm dishes. Cells were transiently treated with siRNA targeting BRAF (5’-AAGUGGCAUGGUGAUGUGGCA-3’) with Lipofectamine RNAiMAX (Thermo Fisher Scientific 13778100). After 24 hours of siRNA treatment, cells were treated with 1μg/mL of doxycycline (Selleck S5159) to turn on the expression of the TurboID constructs. After 48 hours of siRNA treatment and 24 hours of doxycycline treatment, cell culture media was supplemented with 50μM biotin for 30 minutes. Cells were harvested and pellets were washed twice with PBS. Cell pellets were lysed in 700μL of cold RIPA buffer (recipe listed above) with HALT protease inhibitor cocktail (1:50) and incubated on ice for 30 minutes with vortexing every 5 minutes. Lysates were spun at 14,000 x g at 4°C for 10 minutes. 2 mL Zeba® Desalt columns 7k MWCO (Thermo Fisher Scientific) were prepared and supernatant from centrifuged lysates were desalted on the Zeba® columns with a 50μL RIPA stacker. HALT protease inhibitor cocktail was added back to the lysates (1:50) and a BCA was performed to quantify protein concentrations. 1-3mg of equal protein lysate was added to 300 μL of streptavidin dynabeads (Invitrogen 11205D) and were rotated overnight at 4°C. Beads were then collected on a magnetic stand for 1 minute and supernatant was removed. Beads were washed twice with 1mL of RIPA buffer, then 1mL of 1 KCl, followed by 1mL of Na_2_CO_3_ and beads were collected after 10 seconds. Beads were then washed with 1mL of 2M urea in 10mM

Tris HCl pH 8.0 and immediately collected on the magnetic stand. Beads were resuspended in 1mL of RIPA buffer and transferred to fresh tubes. Beads were washed once more with 1mL of RIPA buffer followed by 5 washes in Tris-HCl pH 8.0. For immunoblotting, biotinylated proteins were eluted with 80μL of 2X SDS, DTT, and 2mM biotin and boiled for 5 minutes at 100°C and run as described above. For mass spectrometry analyses, beads were resuspended in 500μL of Tris HCl pH 8.0 and shipped overnight to the Taplin Biological Mass Spectrometry Facility at Harvard University.

### Bead Digestion Analysis by LC-MS/MS

At the Taplin Biological Mass Spectrometry Facility beads were washed at least five times with 100μl 50 mM ammonium bicarbonate then 5μl (200 ng/μl) of modified sequencing-grade trypsin (Promega, Madison, WI) was spiked in and the samples were placed in a 37°C room overnight. The samples were then centrifuged or placed on a magnetic plate if magnetic beads were used and the liquid removed. The extracts were then dried in a speed-vac (∼1 hr). Samples were then re-suspended in 50μL of HPLC solvent A (2.5% acetonitrile, 0.1% formic acid) and desalted by STAGE tip^71^.

On the day of analysis, the samples were reconstituted in 10µl of HPLC solvent A. A nano-scale reverse-phase HPLC capillary column was created by packing 2.6µm C18 spherical silica beads into a fused silica capillary (100 µm inner diameter x ∼30 cm length) with a flame-drawn tip^72^.

After equilibrating the column each sample was loaded via a Famos auto sampler (LC Packings, San Francisco CA) onto the column. A gradient was formed and peptides were eluted with increasing concentrations of solvent B (97.5% acetonitrile, 0.1% formic acid). As peptides eluted they were subjected to electrospray ionization and then entered into a Orbitrap Exploris480 mass spectrometer (Thermo Fisher Scientific, Waltham, MA). Peptides were detected, isolated, and fragmented to produce a tandem mass spectrum of specific fragment ions for each peptide.

Peptide sequences (and hence protein identity) were determined by matching protein databases with the acquired fragmentation pattern by the software program, Sequest (Thermo Fisher Scientific, Waltham, MA)^73^. All databases include a reversed version of all the sequences and the data was filtered to between a one and two percent peptide false discovery rate. Analysis of differentially expressed proteins was performed using DESeq2 (*p*-adjusted <0.05) (PMID: 25516281). Gene Ontology (GO) analysis was performed using the “enrichGo” algorithm in the clusterProfiler package in R (PMID: 22455463).

### Immunofluorescence microscopy

Cells were grown on coverslips to ∼80% confluency. Primary epidermal melanocytes were treated with 200ng/mL of doxycycline for 24 hours. Cells were washed twice with ice cold PBS, fixed at -20°C in methanol for 15 minutes, washed three times with ice cold PBS, and blocked with 10% donkey serum. The primary antibodies used for immunodetected were rabbit-TP53 (Cell Signaling 2527) and mouse-BRAF (Sant Cruz Biotechnology sc-5284). After incubation with fluorescently labeled secondary antibodies anti-mouse 488 (Abcam 150109) and anti-rabbit 594 (Abcam ab150084), coverslips were mounted with DAPI mounting medium (ab1041039) and imaged by Lecia SP8 confocal white light laser (Leica application suite X software) at the University of Utah Cell Imaging Core. Colocalization was quantified on a Lecia confocal microscope using the Leica LasX colocalization analysis (version 3.5.7.23225) software to quantify the overlap between fluorescent signals.

### Proximity ligation assays

Duolink proximity ligation assay (PLA) was performed following the manufacturer’s protocol. Briefly, cells were plated on glass coverslips in 6-well culture plates to ∼80% confluency.

Primary epidermal melanocytes were treated with 200ng/mL of doxycycline for 24 hours. Cells were washed twice with ice cold PBS, fixed at -20°C in methanol for 15 minutes, and washed three times with ice cold PBS. Duolink blocking buffer (Sigma DUO82007) was utilized and rabbit-TP53 1:10,000 (Cell Signaling 2527), rabbit-14-3-3 1:20,000 (Cell Signaling 8312) and mouse-BRAF 1:10,000 (Santa Cruz Biotechnology sc-5284) were used as primary antibodies. After primary antibody incubation, Duolink PLA probes Anti-Mouse MINUS (Sigma DUO92004) and Anti-Rabbit PLUS (Sigma DUO92002) were used as well as the Duolink in situ detection fluroescent green reagent (Sigma DUO92014). Coverslips were mounted onto microscopy slides using DAPI mounting medium (ab1041039). Slides were guarded from light and imaged within 48 hours on a confocal microscope (Lecia SP8 confocal white light laser). Average PLA signals per nucleus were quantified using Leica application suite X software at the University of Utah Cell Imaging Core and ImageJ software (Fiji Version 2.14.0) for a minimum of 3 fields of view per condition.

### Generation of TP53 truncation mutants

The FLAG-TP53 vector was a gift from Dr. Haian Fu (Emory University). Wild-type TP53 was lifted out through restriction digest using BamH1 and PspOMI. TP53 alterations generated through PCR reactions using the following primers **FLAG Forward:** GGTGGATCTGGAGGGTCTGGAGGGGGTGGATCTAAG, **TP53 Reverse:** GGTGACACTATAGAATAGGGCCC. 1)

CGATGACAAGGGTGGTGGATTGATGCTGTCCCCGGAC; 2)

CGATGACAAGGGTGGTGGAGATGAAGCTCCCAGAATGCC; 3)

CGATGACAAGGGTGGTGGACTGTCATCTTCTGTCCCTTCC; 4)

GTGACACTATAGAATAGTCAGGGCCAGGAGGGG; 5)

CGATGACAAGGGTGGTGGAAAAGGGGAGCCTCACCAC; 6)

GTGACACTATAGAATAGTCATCCATCCAGTGGTTTCTTCTTTG; 7) GTGACACTATAGAATAGTCAAGCCTGGGCATCCTTGAG; No DBD) TGGCCCCTGTCATCTTCTAAAGGGGAGCCTCACCAC, GTGGTGAGGCTCCCCTTTAGAAGATGACAGGGGCCAG; DBD) ACGATGACAAGGGTGGTGGATCCCCCCTGTCATCTTCTGTCCCTTC, GGTGACACTATAGAATAGCTATTTCTTGCGGAGATTCTCTTCCTCTG. Digested vector and inserts were combined together at a 5 (insert): 1 (vector) picomolar ratio with Gibson Assembly Master Mix (NEB E2611) and incubated at 50° C for 1 hour. Annealed vectors were transformed into One Shot Mach1 chemically competent cells (Invitrogen C862003) and verified through sanger sequencing.

### Immunoprecipitations

HEK293T cells were seeded in 6-well plates and transiently transfected with FLAG-TP53 constructs and WT or V600E GST-BRAF constructs with Lipofectamine 3000 reagent (Thermo Fisher Scientific L3000015). 48 hours after transfection cells were harvested in ice cold PBS. Washed cells were lysed in an appropriate amount of lysis buffer 400 (50mM Tris pH 8.0, 400mM NaCl, 1% Nonidet P-40 substitute, 10% glycerol) and HALT protease inhibitor cocktail. Beads were incubated with 1μg of FLAG-antibody for 1 hour with rotation at 4°C and then washed three times with 1X TBS with Tween-20. A BCA was performed to quantify protein concentrations. 300-500 μg of equal protein lysate was added to 50μL of magnetic Dynabeads® (Thermo Fisher Scientific 10004D) and rotated at 4°C for one hour. Beads were washed three times with 1X TBS and immunoprecipitates were eluted with NuPAGE LDS Sample Buffer (Thermo Fisher Scientific NP0007) and NuPAGE Sample Reducing Agent (Thermo Fisher Scientific NP0009) and immunoblots were run as described above.

### Nutlin-3a treatment

Nutlin-3a was obtained from Sellek chem and was resuspended in DMSO. Cells were treated with varying concentrations of Nutlin-3a (0-20μM) for 24 hours before cell lysates were harvested for immunoblotting.

### RNA Sequencing

RNA was collected from 2 biological replicates of the primary epidermal melanocyte cells treated with or without 200ng/mL of doxycycline. Cells were collected directly into Trizol and stored at - 80°C until purification. RNA was isolated via Trizol-chloroform extraction followed by column- based purification. The aqueous phase was brought to a final concentration of 35% ethanol, and RNA was purified using the PureLink RNA Mini kit according to the manufacturer’s instructions (ThermoFisher Scientific). Library preparation was performed using the NEBNext Ultra II Directional RNA Library Prep with poly(A) mRNA isolation. Sequencing was performed using the Illumina NovaSeq 6000 (150 x 150 bp paired-end sequencing, 25 million reads per sample).

### RNA-seq data processing and analysis

The human hg38 genome and gene feature files were downloaded from Ensembl and a reference database was created using STAR version 2.7.6a^74^. Optical duplicates were removed from NovaSeq runs via Clumpify v38.34^75^. Reads were trimmed of adapters and aligned to the reference database using STAR in two-pass mode to output a BAM file sorted by coordinates. Mapped reads were assigned to annotated genes using featureCounts version 1.6.3^76^. Raw counts were filtered to remove features with zero counts and features with five or fewer reads in every sample. DEGs were identified using the hciR package (https://github.com/HuntsmanCancerInstitute/hciR) with a 5% false discovery rate and DESeq2 version 1.34.0^77^.

### Dual luciferase assays

Cell lines were transiently transfected with PG13-luc (wt TP53 binding sites), a gift from Bert Vogelstein (Addgene plasmid # 16442 ; http://n2t.net/addgene:16442 ; RRID:Addgene_16442), pRL-CMV-Renilla control vector (Promega E2261), and lipofectamine 3000 (Thermo Fisher Scientific L3000150). Promega Dual Luciferase Assay was performed based on the manufacturer’s protocol.

### Analysis of human melanoma **-** TCGA

We filtered all TCGA Pan Cancer Atlas cutaneous melanoma samples based on TP53 mutation status. Kaplan Meyer survival curves were generated using R. Survival analysis was performed using a Cox proportional hazards model. Differences in survival between groups is shown using hazard ratios and log-rank test p-values.

### Analysis of BRAF mutations across cancers **-** cBioPortal

We utilized cBioPortal and the Onco Query Language (OQL) to query for patients with BRAFV600E mutations and any *TP53* alteration to determine mutual exclusivity or co-occurrence in cancers commonly driven by oncogenic BRAF including cutaneous melanoma, lung adenocarcinoma, colorectal carcinoma, and thyroid cancer. Low p-values indicate a statistically significant tendency towards mutual exclusivity based on a one-sided Fisher’s Exact Test. The p-values are listed in the table.

### Nuclear: Cytoplasmic Fractionation

Cellular fractionation experiments were performed using the NE-PER Nuclear and Cytoplasmic Extraction Kit (Thermo 78833). Briefly, cells pellets were collected by centrifugation at 1,000 xg for 1 minute. Cell pellets were then resuspended in an appropriate volume of CER I buffer based on the packed cell volume of each pellet. Cells were vortexed and incubated on ice for 10 minutes. CER II buffer was added to each tube, and samples were briefly vortexed prior to a 1- minute incubation on ice. Cell lysate was then pelleted at max speed for 5 minutes and the supernatant (cytoplasmic fraction) was transfer to a new, pre-chilled 1.5mL tube. The remaining insoluble pellet (nuclear fraction) was resuspended in the appropriate amount of ice-cold NER buffer and vortex for 15 seconds every 10 minutes for a total of 40 minute incubation on ice. Nuclear lysate was then pelleted at max speed and the supernatant was transferred to a new 1.5mL tube. Cell lysate was quantified using a BSA assay.

### Alphafold

To visualize the *in vivo* protein-protein interactions between BRAF and TP53, a prediction model was run on AlphaFold3 (DeepMind, London, UK). Amino acid sequences of BRAF and TP53 were obtained from the Universal Protein Resource website (UniProt.org) using the identifiers P15056 and P04637. These sequences were then analyzed together using the AlphaFold3 server. The highest ranked predictions were then modeled in ChimeraX software as indicated in the figures. Interface diagrams of the modeled BRAF-TP53 complexes were generated in ChimeraX by using the interface command found within the molecule display tab. The interface surface area is determined by ChimeraX MSMS modeling (surface modeling) wherein a probe with the radius of a water molecule (1.4 Å) is used to calculate the solvent-accessible surface area of a molecule.

### Quantification and statistical analysis

All graphing and statistical analysis was performed with PRISM software or R, with all graphs showing mean and standard deviation or standard error. The statistical details can be found in the corresponding figure legend. All NGS statistical analysis was performed according to published pipeline protocols cited, with a statistical significance cutoff of padj<0.05.

## Acknowledgements

We are grateful to acknowledge: 1. Members of the McMahon lab for their helpful suggestions and comments; 2. Dr. Sheri Holmen for reagents and suggestions. 3. Dr. Martin Golkowski for helpful suggestions on mass spectrometry experiments; 4. Dr. Doug Grossman for advice, guidance, and access to a UV exposure chamber; 5. Dr. Wes Sundquist for proximity labeling expertise and guidance; 6. Dr. Ben Myers for guidance on AlphaFold and structural studies; 7. The HSC DNA sequencing core for assistance with Sanger sequencing and cell line authentication; 8. The HSC Flow Cytometry Core Facility for assistance with single cell sorting; 9. The Cell Imaging Core Facility for assistance with confocal microscopy; 10. Ross Tomaino and the Taplin Mass Spectrometry Facility at Harvard University for assistance with the mass spectrometry for the TurboID screen. The results published here are in part based upon data generated by the TCGA Research Network: https://www.cancer.gov/tcga. M.M. was supported by grants from the NIH (R01CA176829 and R01CA131261) and institutional funds from the Melanoma Disease Center and the HCI Comprehensive Cancer Support grant (P30 CA042014). K.T.O. was supported by the NIH/NCI (T32CA265782). K.T.O. was also supported by the Huntsman Cancer Foundation. E.L.S. was supported by grants from the NIH (R01CA212415, R01CA240317 and R01CA237404). G.F. was supported by the NIH/NCI (F31CA275328). R.J.T. and A.P. were supported by the NIH/NCI (R01CA229896). The content is solely the responsibility of the authors and does not necessarily represent the official views of the NIH.

## RESOURCE AVAILABILITY

### Lead contact

Further information and requests for resources and reagents should be directed to and will be fulfilled by the lead contact, Martin McMahon

### Materials availability

TurboID constructs generated in this study are available upon request.

### Data availability

All data generated or analyzed during this study are included in the manuscript and supporting files; source data files have been provided for Figure 1 and supplemental figure 1.

---

**Supplemental Figure 1:**
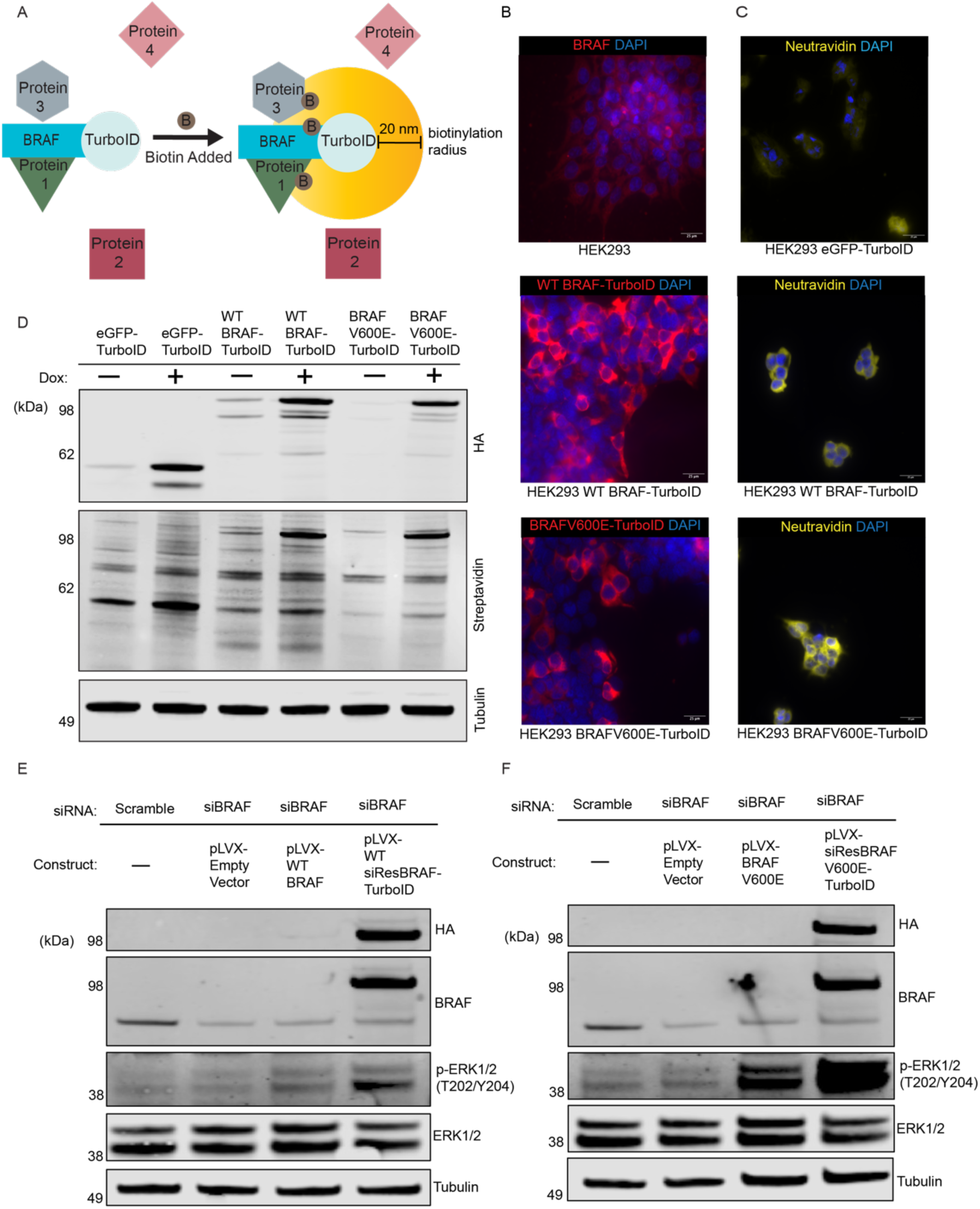
Characterization and validation of TurboID constructs. **A.** Schematic depicting BRAF-TurboID fusion proteins and inherent ability to biotinylate proximal protein interactors, such as proteins 1 and 3, but not proteins 2 and 4. **B.** Representative images of endogenous BRAF and transiently transfected HA-BRAF-TurboID (wild-type and V600E) constructs in HEK293 cells as visualized by fluorescence microscopy with primary antibodies to BRAF and HA, respectively. (Scale bar = 25μM. Representative images shown from n=3 biological replicates). **C.** Representative images of TurboID constructs with biotinylation of endogenous proteins visualized by fluorescent neutravidin. (Scale bar = 25μM. Representative images shown from n=3 biological replicates). **D.** Immunoblot analyses demonstrate that all TurboID constructs are expressed following 1 μg/mL of doxycycline treatment, resulting in elevated protein biotinylation. **E.** Immunoblots depicting transient expression of a series of plasmids including empty vector, wild-type BRAF, and siRNA-resistant wild-type BRAF-TurboID with and without siBRAF treatment. **F.** Immunoblots depicting transient expression of a series of plasmids including empty vector, BRAF^V600E^, and siRNA-resistant BRAF^V600E^-TurboID with and without siBRAF treatment.

